# A Synergistic Terpenoid–Phenolic Defense Network Confers Resistance to *Dendrolimus superans* in Hybrid Larch

**DOI:** 10.64898/2026.10.07.757267

**Authors:** Yang Panke, Liu Maolan, Xin Linhuai, Liu Jing, Li Chunming, Bai Hui

## Abstract

To elucidate the resistance of hybrid larch (*Larix* spp.) clones to *Dendrolimus superans* and their chemical defense mechanisms, 46 clones were subjected to field feeding–infestation experiments. Insect resistance was comprehensively evaluated by the membership function method from three perspectives: insect weight change, frass weight and defoliation rate (damage consequences); pre- and post-infestation physiological indices (constitutive and induced resistance). High- and low-resistance materials screened from each perspective were further analyzed by widely targeted LC-MS metabolomics. The three damage indices differed highly significantly among clones (*P*<0.01) and were positively correlated, indicating abundant genetic variation in resistance. Clones such as -30-C, -35-A and -21-A showed strong overall resistance, whereas -28-E and -20-A showed high constitutive and induced resistance. Defence enzyme activities (POD, SOD, CAT, PPO and PAL) correlated closely with secondary metabolites (total phenolics, tannins and total flavonoids) and nutrients, and rose synergistically after feeding, reflecting synergy between constitutive and induced resistance. In total, 3,577 metabolites were identified, predominantly terpenoids (>90%). The three groups yielded 1,010, 1,086 and 922 differentially accumulated metabolites, and PCA separated high-from low-resistance materials in all groups. These metabolites were significantly enriched in metabolic pathways, biosynthesis of secondary metabolites, monoterpenoid and diterpenoid biosynthesis, and phenylpropanoid biosynthesis. In conclusion, resistance of hybrid larch to *D. superans* is jointly conferred by the defence enzyme system, secondary metabolite accumulation and a “terpenoid–phenolic” metabolic network: terpenoids and phenolics act synergistically via the terpenoid and phenylpropanoid pathways, together forming the core of chemical defense and supporting the breeding of insect-resistant clones.

## Introduction

Larch (*Larix* spp.) grows rapidly with excellent wood quality and strong adaptability, and is an important timber and ecological forest species in northeastern China^1^. Hybrid larch has been widely promoted for its fast growth, high yield and adaptability^2,3^. However, the *Dendrolimus superans* is the most important defoliating pest of larch plantations in this region; its larvae feed on needles, causing sharp growth reduction and even large-scale defoliation and tree mortality^4^, and chemical control after outbreaks is costly, environmentally problematic and increasingly ineffective due to insecticide resistance^5^. Selecting insect-resistant superior materials is therefore the most economical and sustainable management strategy. Yet insect resistance breeding faces a core trade-off: resources allocated to defense are drawn from the carbon and nitrogen reserves required for growth, so enhanced resistance often reduces growth^1,6^. Therefore, how to screen out superior materials that can resist pests without significantly compromising growth remains a key issue that urgently needs to be addressed in forest insect resistance breeding.

Plant resistance to defoliating pests is not a single trait but has a distinct two-dimensional nature: constitutive resistance, present before feeding^7,8^, and induced resistance, activated in response to pest feeding^9,10^. The former is determined by pre-existing defense enzyme activity, secondary metabolite reserves and nutrient levels, forming the first line of defense^11,12^; the latter is amplified under feeding stress through activation of the defense enzyme system and rapid synthesis of secondary metabolites, representing the dynamic response to attack^13,14^. These two forms are driven by distinct mechanisms and wax and wane complementarily across stress stages, jointly determining final resistance^7,15^. However, existing studies on larch insect resistance have mostly focused on a single index at one hierarchical level or time point, evaluating constitutive and induced resistance in isolation, which cannot comprehensively reflect the true resistance level of a material^16,17^.

Conifers are rich in terpenoids and phenolics, which together constitute the core of their chemical defense. Terpenoids, such as diterpene resin acids, monoterpene volatiles and iridoid glycosides, generally possess direct antifeedant and toxic activities interfering with pest feeding, digestion and development^18,19^. Phenolics (e.g., phenylpropanoids)not only participate in cell wall reinforcement as lignin precursors to form a physical barrier, but also exhibit insecticidal activity and can serve as precursors of phytoalexins^20,21^. Terpenoids and phenolics thus do not act independently but synergistically through the terpenoid and phenylpropanoid metabolic pathways, forming a multi-layered, inducible chemical defense network^22,23^.

Accordingly, this study used 46 hybrid larch clones in field feeding–infestation experiments with *D. superans*, comprehensively evaluating insect resistance by the membership function method from three perspectives: insect weight, frass weight and defoliation rate (damage consequences); pre-infestation physiological indices (constitutive resistance); and post-infestation physiological indices (induced resistance). Representative high-and low-resistance materials were screened and subjected to widely targeted LC-MS metabolomics. The objectives were to (1) clarify the pattern of genetic variation in insect resistance and the multi-perspective evaluation results, providing a basis for screening superior materials; (2) reveal at the metabolomic level the key differentially accumulated metabolites and pathways through which high- and low-resistance materials respond to feeding; and (3) elucidate the synergistic “terpenoid–phenolic” chemical defense network, corroborated by physiological indices before and after feeding. This systematically interprets the resistance mechanism at the phenotypic, physiological and metabolic levels, providing a theoretical basis and material foundation for breeding insect-resistant superior clones.

## 2 Materials and Methods

### 2.1 Plant Materials

The 46 hybrid larch clones were obtained from Qingshan Forest Farm, Linkou County, Mudanjiang City, Heilongjiang Province, China (133°53′E, 46°38′N; Table 1). They were grafted onto *Larix olgensis* rootstocks in May 2024 and, after one year of successful survival, were lifted and transported in April 2025 to Maosen Nursery Professional Planting Cooperative, Yangming District, Mudanjiang City, Heilongjiang Province (129°26′E, 44°56′N), where they were planted in 20 × 20 cm seedling pots.

**Table 1.** Information of Tested Hybrid Larch Clones.

| No. | Clone ID | No. | Clone ID |
| --- | --- | --- | --- |
| 1 | □-26-C | 24 | □-14-A |
| 2 | □-25-D | 25 | □-31-B |
| 3 | □-31-A | 26 | □-31-C |
| 4 | □-10-A | 27 | □-23-A |
| 5 | □-26-B | 28 | □-4-C |
| 6 | □-7-A | 29 | □-21-B |
| 7 | □-35-A | 30 | □-4-A |
| 8 | □-26-A | 31 | □-22-A |
| 9 | □-30-C | 32 | □-20-A |
| 10 | □-15-A | 33 | □-28-E |
| 11 | □-21-A | 34 | □-1-A |
| 12 | □-16-A | 35 | □-5-A |
| 13 | □-12-C | 36 | □-10-7 |
| 14 | □-31-A | 37 | 5-5-1 |
| 15 | □-19-A | 38 | 2-10-6 |
| 16 | □-10-C | 39 | 25-16-17 |
| 17 | □-21-A | 40 | 35-6-5 |
| 18 | □-32-A | 41 | 5-10-11 |
| 19 | □-8-A | 42 | 6-2-6 |
| 20 | □-25-A | 43 | 7-4-17 |
| 21 | □-7-A | 44 | 6-5-4 |
| 22 | □-6-C | 45 | 37-1-21 |
| 23 | □-10-A | 46 | 6-5-15 |

On 10 May 2025, third-instar *D. superans* larvae were collected in Hongmei Town, Meihekou City, Jilin Province, China (125°59′E, 42°41′N) and reared under natural conditions at the experimental site (129°26′E, 44°56′N). Each acrylic rearing box (15 × 20 × 30 cm) contained 30 larvae fed with fresh larch needles replaced every other day, with residual debris and frass removed. Field feeding–infestation began once the larvae had molted to the fourth instar.

### 2.3 Field Infestation Experiment

The experiment commenced in mid-to-late May 2025. Beforehand, larvae were starved for 24 h to eliminate residual gut contents. Five randomly selected larvae were placed in a Petri dish, weighed (0.001 g precision), then transferred into a 40-mesh gauze bag (40 × 50 cm) placed over a clone ramet seedling; five ramets were randomly selected per clone, and the bag was tied tightly at the graft union to prevent escape or feeding on rootstock needles. After 72 h, bags were removed, frass collected, and larvae reweighed. Frass was oven-dried at 80 °C for 12 h and weighed. Needles from the scion portion were then collected, transported on dry ice, and stored at −80 °C.

### 2.4 Determination of Physiological and Biochemical Indices

For each clone, five ramets were randomly selected and newly developed scion needles were collected 3 days before the experiment, transported on dry ice, and stored at −80 °C. Using reagent kits (Suzhou Geruisi Biotechnology Co., Ltd., China), the following were determined in needles collected before and after feeding: peroxidase (POD, G0107W), superoxide dismutase (SOD, G0101W), catalase (CAT, G0105W), polyphenol oxidase (PPO, G0113W), phenylalanine ammonia-lyase (PAL, G0114W), tannin (G0119W), total phenolics (TP, G0117W), total flavonoids (TF, G0118W), amino acid (AA, G0415W), soluble protein (SP, G0417W, Coomassie brilliant blue method) and soluble sugar (SS, G0501W) contents.

### 2.5 Related Calculation Formulas

## (1) Insect weight change

The formula for insect weight change is as follows: *W* = *W*2-*W*1

Where: *W*2 is the weight of the *D. superans* larvae after feeding, and *W*1 is the weight of the larvae before feeding.

## (2) Defoliation rate

Defoliation rate refers to the degree of needle loss in the trees, usually expressed as the ratio of the amount of canopy foliage lost to the total amount of foliage per unit area. The defoliation rate of the hybrid larch clones was calculated by reference to the method of Saheya^24^.

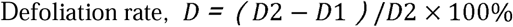

Where: *D*2 is the total number of needles before feeding by the *D. superans*, and *D*1 is the number of remaining needles after feeding.

1. The insect resistance of the tested materials was comprehensively evaluated using the membership function method, and the calculation formula is as follows^25^:

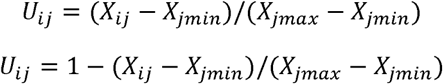

Where: *U* is the membership function value of the i-th sample for the j-th index; X*_ij_* is the measured value of the j-th index for the i-th sample; and *X_jmin_ X_jmax_* are the minimum and maximum values of the j-th index, 1,2,3… … *n*

The entropy weight method was used for weighting, and the entropy value of the j-th index was calculated as follows:

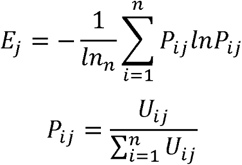

Where:*n* is the number of samples.

The difference coefficient of the j-th index was calculated as follows:

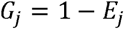

The weight of the j-th index was calculated as follows:

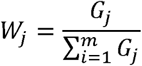

Where: m is the number of evaluation indices measured.

The comprehensive index was calculated as follows:

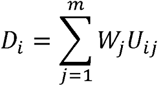

Where: D is the comprehensive index of the i-th sample; W is the weight of the j-th index; and U is the membership function value of the i-th sample for the j-th index. The value of D reflects insect resistance, with a higher value indicating greater insect resistance.

### 2.6 Metabolomic Analysis

Based on the comprehensive evaluation, 6 groups comprising 18 samples (3 biological replicates each) were selected for LC-MS metabolomic analysis. For metabolite extraction, samples were freeze-dried and ground (60 Hz, 30 s); 25 mg (±1 mg) was transferred to a 2 mL tube with 1000 µL extraction solvent (methanol:water, 4:1, v/v, containing an internal standard); two steel balls (3.2 mm) were added, vortexed for 30 s, homogenized at 45 Hz for 4 min and ultrasonicated in an ice-water bath for 5 min, repeated three times; samples were then held at −40 °C for 1 h and centrifuged at 4 °C, 12,000 rpm for 15 min, and the supernatant was filtered (0.22 µm) for analysis (reagents and instruments in Tables 2 and 3). UHPLC separation used a Vanquish system (Thermo Fisher Scientific) with mobile phase A (0.01% acetic acid in water) and B (50% acetonitrile/isopropanol); column temperature 25 °C, autosampler 4 °C, injection volume 2 µL, flow rate 0.3 mL/min. Detection used a Stellar mass spectrometer (Thermo Fisher Scientific) with spray voltage 3.8 kV/−3.4 kV, sheath gas 50 Arb, aux gas 15 Arb, sweep gas 1 Arb, ion transfer tube 320 °C and vaporizer 450 °C. Differential metabolites were selected using |log Fold Change| > 1 and PADJ < 0.05, further screened by VIP > 1 from the OPLS-DA model, with significance assessed by a two-tailed Student’s t-test (P < 0.05). Data acquisition and quantification used Biobud software (Biotree Biotech Co., Ltd., China).

**Table 2.** Reagents for Metabolomics Analysis.

| Chemical Name | CAS | Purity | Source |
| --- | --- | --- | --- |
| Methanol | 67-56-1 | LC-MS | CNWTechnologies (Shanghai Bailey Biotechnology Co., Ltd.) |
| Acetonitrile | 75-05-8 | LC-MS | CNWTechnologies (Shanghai Bailey Biotechnology Co., Ltd.) |
| ddH<sub>2</sub>O | - | - | Watsons (Guangzhou, Guangdong Province) |
| Acetic acid | 64-19-7 | LC-MS | SIGMA-ALDRICH(Sigma-Aldrich (Shanghai) Trading Co., Ltd.) |
| 2-Propanol | 67-63-0 | LC-MS | CNWTechnologies (Shanghai Bailey Biotechnology Co., Ltd.) |

**Table 3.** Instruments for Metabolomics Analysis.

| Instrument | Model | Manufacturer |
| --- | --- | --- |
| UHPLC | Vanquish | Thermo Fisher Scientific (Massachusetts, USA) |
| HRMS | Orbitrap Exploris 120 | Thermo Fisher Scientific (Massachusetts, USA) |
| Centrifuge | Heraeus Fresco17 | Thermo Fisher Scientific (Massachusetts, USA) |
| Analytical balance | BSA124S-CW | Sartorius (Göttingen, Germany) |
| Ultrasonic bath | PS-60AL | Shenzhen Leidebang Electronic Co., Ltd. |
| Homogenizer | JXFSTPRP-24 | Shanghai Jingxin Technology Co., Ltd. |
| Freeze dryer | LGJ-10C | Sihuan Furuikeyi Technology Development Co., Ltd. |

Metabolite extraction (the reagents and instruments used are shown in Tables 2 and 3): (1) The samples were freeze-dried and then ground (60 Hz, 30 s). (2) A precisely weighed amount of 25 mg (±1 mg) of each sample was transferred into a 2 mL centrifuge tube, and 1000 μL of extraction solvent (methanol: water, 4:1, v/v, containing an internal standard) was added. (3) Two steel balls (3.2 mm) were added, and the mixture was vortexed for 30 s, homogenized at 45 Hz for 4 min, and ultrasonicated in an ice-water bath for 5 min. (4) The homogenization and ultrasonication were repeated three times. (5) The samples were placed in a −40 °C refrigerator and allowed to stand for 1 h. (6) The samples were centrifuged at 4 °C and 12,000 rpm for 15 min, and the supernatant was filtered through a 0.22 μm membrane into an injection vial for instrumental analysis.

LC-MS/MS analysis: UHPLC separation was carried out using a Vanquish system (Thermo Fisher Scientific). Mobile phase A was 0.01% acetic acid in water, and mobile phase B was 50% acetonitrile/isopropanol (ACN/IPA). The column temperature was set at 25 °C and the autosampler temperature at 4 °C; the injection volume was 2 μL and the flow rate was 0.3 mL/min. A Stellar mass spectrometer (Thermo Fisher Scientific) was used for sample detection. The ion source parameters were as follows: spray voltage, 3.8 kV/−3.4 kV; sheath gas, 50 Arb; aux gas, 15 Arb; sweep gas, 1 Arb; ion transfer tube temperature, 320 °C; and vaporizer temperature, 450 °C. Differential metabolites were selected using |log Fold Change| > 1 and PADJ < 0.05 as thresholds and further screened by a variable importance in projection (VIP) value > 1 derived from the OPLS-DA model. To assess the significance of differences in metabolite abundance, a two-tailed Student’s t-test was used (*P*<0.05).

All mass spectrometry data acquisition and quantitative analysis of the target compounds were performed using the Biobud automated data processing software developed by Biotree Biotech Co., Ltd.

### 2.7 Data Processing and Statistical Analysis

Data organization and formula calculation used Microsoft Excel 2024; analysis of variance, multiple comparisons and Pearson correlation analysis used SPSS 27.0; hierarchical cluster analysis and figure generation used Origin 2024 Pro.

## 3 Results

### 3.1 Differences in Field Feeding–Infestation of the *D. superans* among Clones

The results of the statistical analysis of insect weight change, frass weight and clone defoliation rate following feeding–infestation by the *D. superans* on different hybrid larch clones are shown in Table 4. As shown in the table, insect weight change, frass weight and defoliation rate all differed highly significantly among the clones fed upon by the *D. superans* (*P* < 0.01). Among them, the insect weight change was highest for the clone -14-A, whereas both frass weight and defoliation rate were highest for the clone -21-B (Table S1).

**Table 4.** Analysis of Differences in Insect Development and Defoliation Rate among Clones.

| Traits | Mean | Range | SD | *F* | *P* |
| --- | --- | --- | --- | --- | --- |
| Insect Weight | 0.70 | 0.17~1.54 | 0.03 | 12.62 | 0.01 |
| Frass Weight | 0.20 | 0.01~0.46 | 0.01 | 4.011 | 0.01 |
| Defoliation Rate | 0.13 | 0.02~0.98 | 0.01 | 13.819 | 0.01 |

Note: $P < 0.05$ indicates significant difference, and $P < 0.01$ indicates highly significant difference.

The results of the correlation analysis between larval development and the phenotypic traits of the clones after the *D. superans* fed on different hybrid larch clones are shown in Fig. 1. As can be seen from the figure, insect weight was significantly positively correlated with defoliation rate (*P* < 0.05), and frass weight was highly significantly positively correlated with defoliation rate (*P* < 0.01).

**Figure 1.**
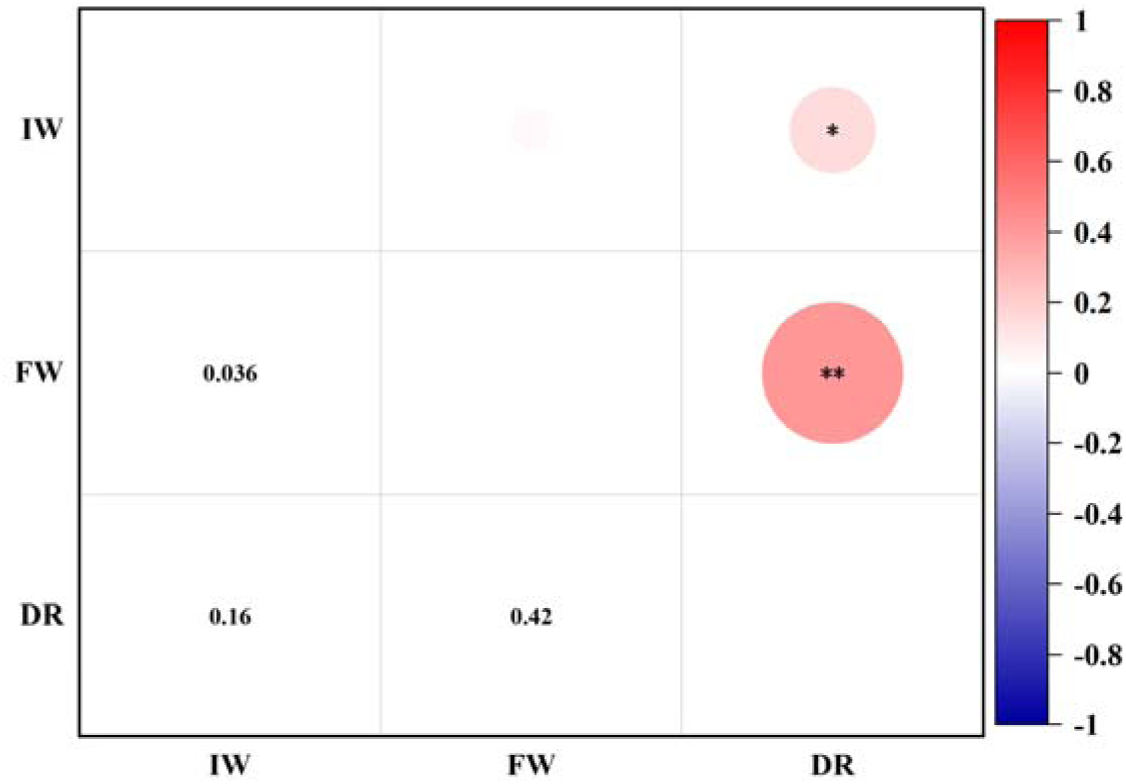
Correlation Analysis between Insect Development and Defoliation Rate of Clones Note: *\** indicates significant difference (*P* < 0.05), and **\*\*** indicates highly significant difference (*P* < 0.01). IW, insect weight; FW, frass weight; DR, defoliation Rate.

Hierarchical cluster analysis of the 46 clones based on these three indices (Fig. 2) separated them into three categories at a genetic distance of 0.8: Category I (31 clones) with insect weight and defoliation rate below the overall mean; Category II (14 clones) with insect weight above the mean; and Category III (one clone) with frass weight and defoliation rate above the mean. Category I thus showed better resistance than Categories II and III. These indices stably reflect feeding damage and reveal abundant genetic variation, and can therefore serve as important phenotypic indicators for evaluating insect resistance.

**Figure 2.**
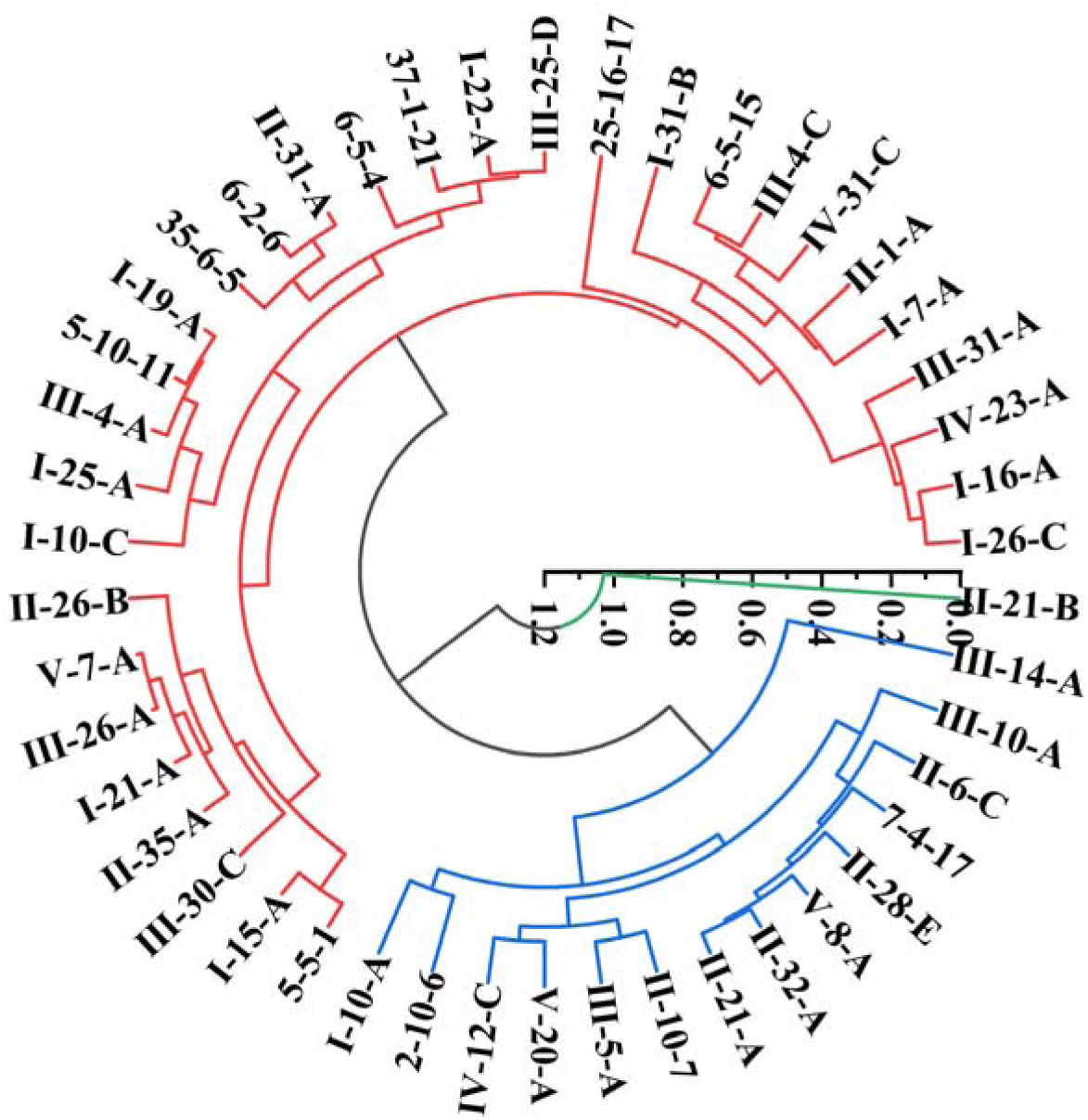
Cluster Analysis of Insect Development and Defoliation Rate of Clones

### 3.2 Differences in Resistance-related Physiological Indices of the Clones before Feeding–infestation by the *D. superans*

Statistical analysis of the resistance-related physiological and biochemical indices of the 46 hybrid larch clones revealed highly significant differences among different clones (*P* < 0.01) (Table 5). Specifically, the POD activity of 5-5-1 was significantly higher than that of the other clones; the SOD activity of 5-10-11 was the highest; the CAT activity of -21-B was the highest; the PPO activity of -26-B was the highest; the PAL activity of 5-5-1 was the highest; the total phenolic content of 6-5-15 was the highest; the tannin content of -28-E was the highest; the total flavonoid content of -5-A was the highest; the amino acid content of -21-A was the highest; the soluble protein content of -26-C was the highest; and the soluble sugar content of -12-C was the highest (Table S2).

**Table 5.**
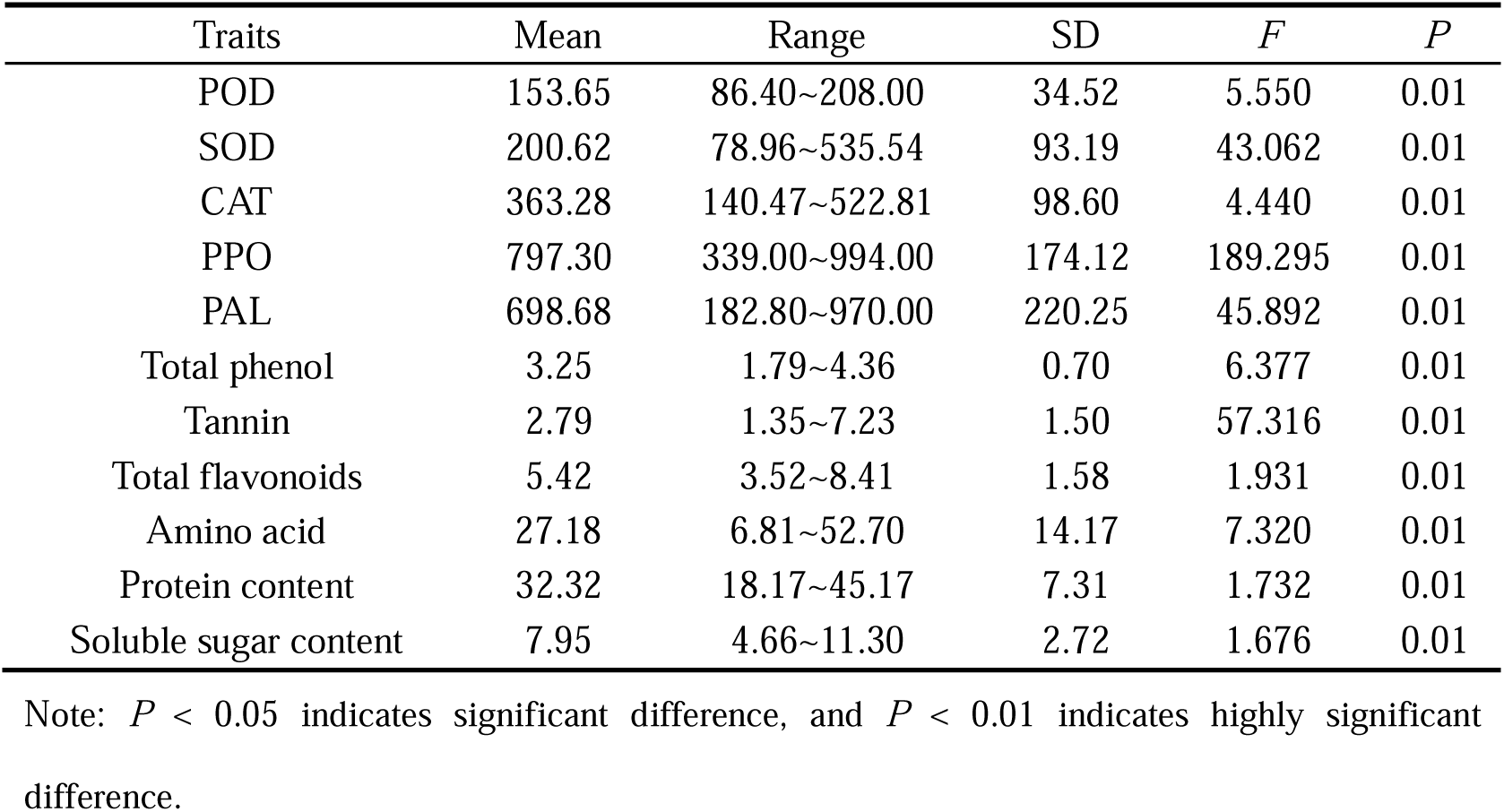
Differences in Physiological Resistance Indices among Clones prior to Feeding by *D. superans*.

Pearson correlation analysis (Fig. 3) showed that POD and SOD activities were highly significantly positively correlated (*P* < 0.01), indicating possible synergy in scavenging reactive oxygen species (ROS), whereas SOD was highly significantly negatively correlated with both PPO and PAL activities (*P* < 0.01), possibly reflecting functional differentiation or substrate competition among enzyme systems during early stress response. PPO and PAL were highly significantly positively correlated (*P* < 0.01), suggesting a close regulatory link between polyphenol oxidase and key phenylpropanoid pathway enzymes. Total phenolics, tannins and total flavonoids were significantly or highly significantly positively correlated with one another, indicating pronounced synergy in secondary metabolite accumulation that may jointly form a chemical defense barrier. These secondary metabolites were significantly negatively correlated with primary metabolites such as amino acids and sugars, reflecting possible competition for carbon or nitrogen between secondary metabolite synthesis and primary metabolism. In summary, before feeding the enzymatic defense, secondary metabolites and nutrients of the clones were highly interrelated, with synergistic secondary metabolite accumulation antagonistic to amino acid nutrients; this interaction network may importantly influence subsequent defense responses.

**Figure 3.**
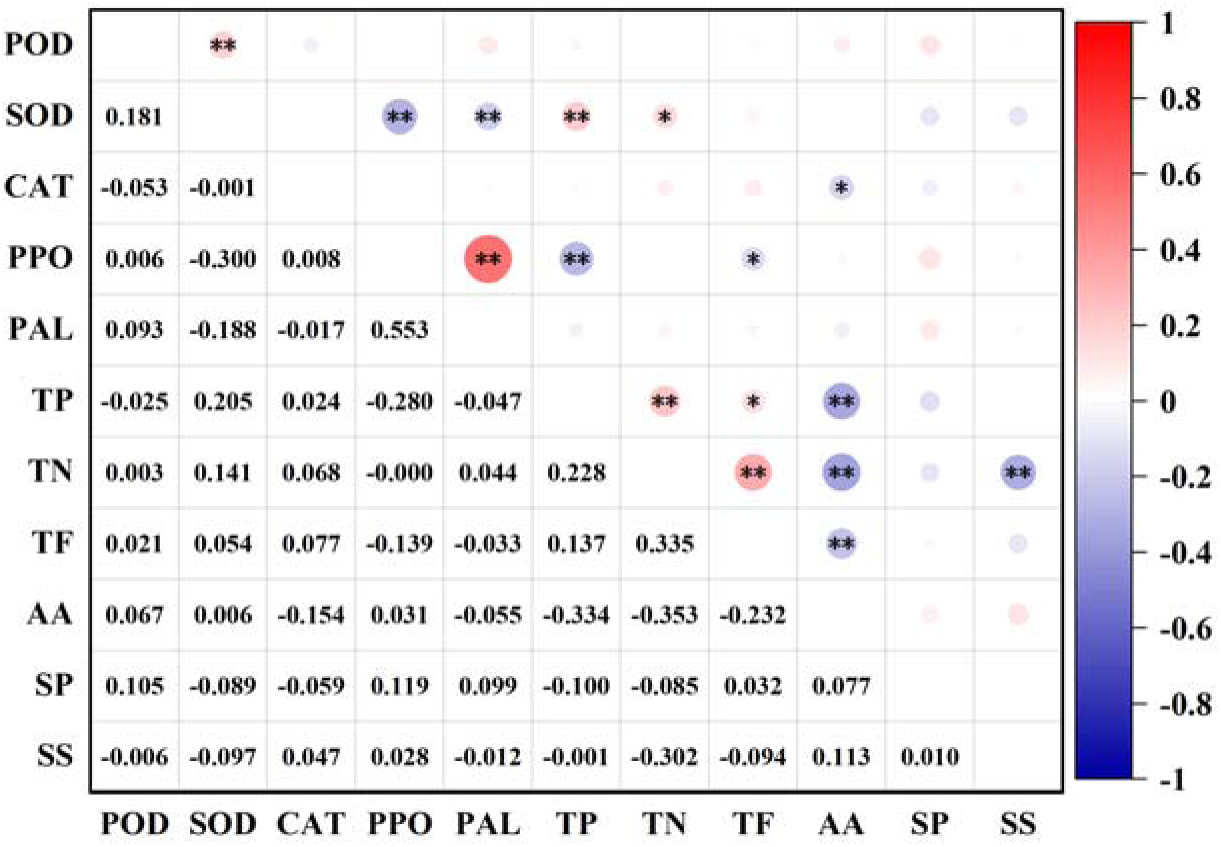
Correlation Analysis of Physiological Resistance Indices among Clones prior to Feeding by *D. superans* Note: *\** indicates significant difference (*P*<0.05), and **\*\*** indicates highly significant difference (*P*<0.01). TP, total phenols; TN, tannins; TF, total flavonoids; AA, amino acids; SP, soluble protein; SS, soluble sugar.

Cluster analysis based on pre-infestation indices (Fig. 4) separated the 46 clones at a genetic distance of 450 into three categories: Category I (36 clones) with relatively higher POD, PPO and PAL activities and tannin, amino acid, soluble protein and soluble sugar contents; Category II (7 clones) with relatively higher POD and SOD activities and total phenolic, total flavonoid and soluble sugar contents; and Category III (3 clones) with relatively higher SOD, CAT and PPO activities and amino acid content. A highly interrelated physiological interaction network had thus already been established before feeding, and this constitutive resistance basis may importantly influence subsequent resistance to feeding.

**Figure 4.**
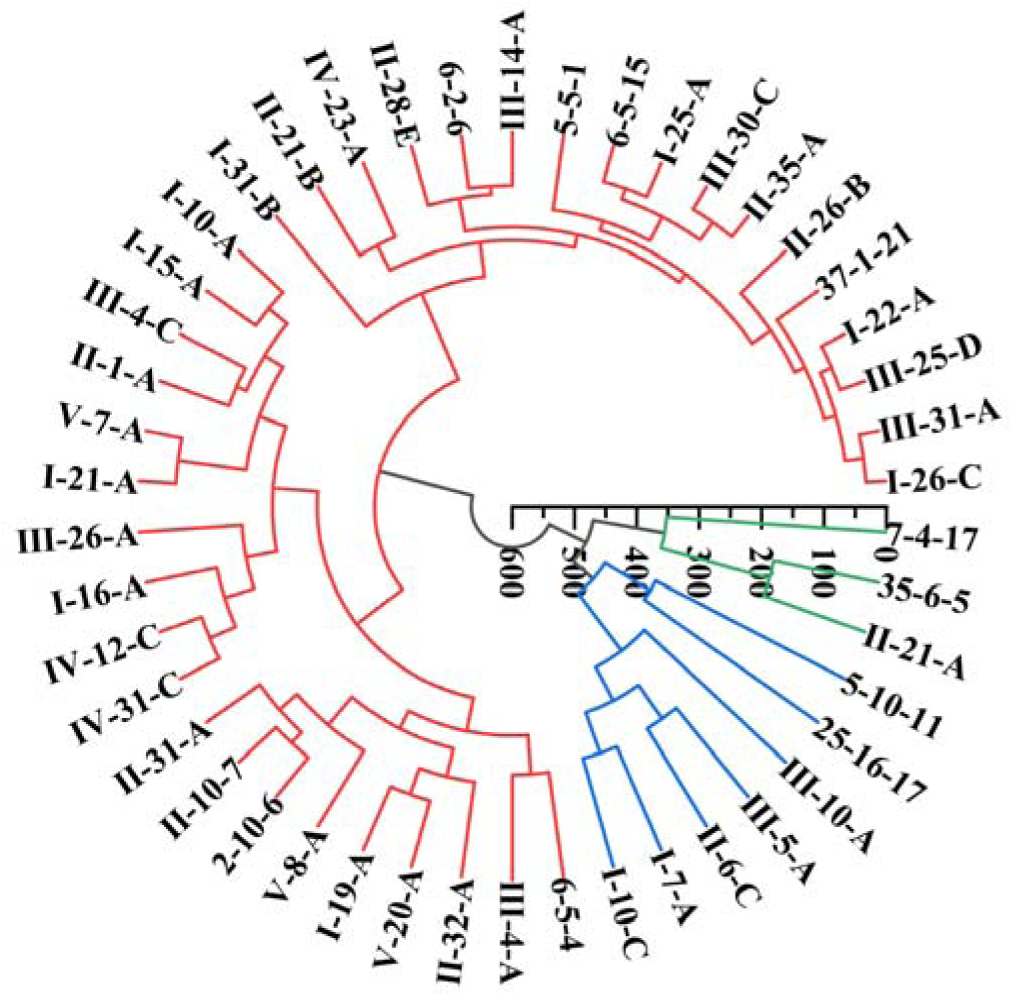
Cluster Analysis of Clones before Infestation by *D. superans*

### 3.3 Differences in Resistance-related Physiological Indices of the Clones after Feeding–infestation by the *D. superans*

Statistical analysis of the resistance-related physiological indices of the 46 hybrid larch clones after feeding–infestation by the *D. superans* revealed highly significant differences among different clones (*P* < 0.01) (Table 6). After feeding–infestation, the POD activity of -19-A was higher than that of the other clones; the SOD activity of 5-10-11 was the highest; the CAT activity of -20-A was the highest; the PPO activity of -26-B was the highest; the PAL activity of -30-C was the highest; the total phenolic content of 6-5-15 was the highest; the tannin content of -20-A was the highest; the total flavonoid content of -5-A was the highest; the amino acid content of -30-C was the highest; the soluble protein content of -25-D was the highest; and the soluble sugar content of -6-C was the highest (Table S3).

**Table 6.**
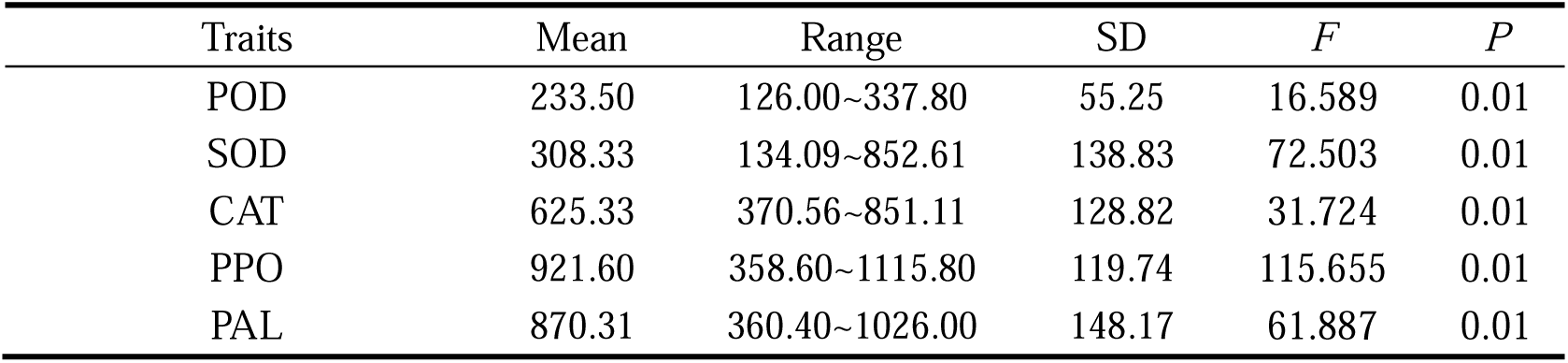
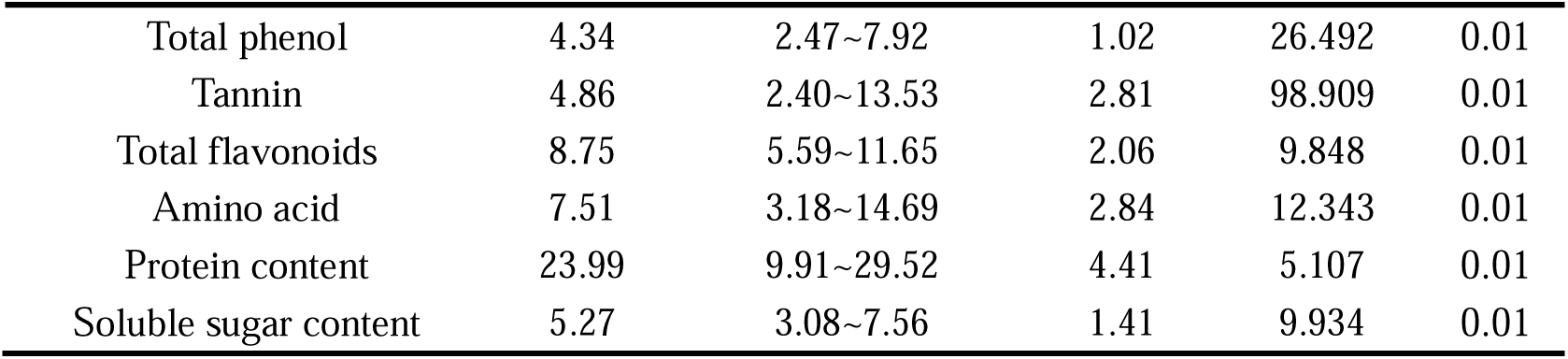
Analysis of Differences in Resistance Physiological Indicators of Clones after Infestation by *D. superans*.

Pearson correlation analysis (Fig. 5) showed that POD was highly significantly positively correlated with both SOD and CAT (*P* < 0.01), indicating synergy among POD, SOD and CAT in scavenging ROS under pest stress. PPO and PAL were highly significantly positively correlated (*P* < 0.01), indicating a close link between key phenylpropanoid pathway enzymes in jointly participating in stress defense. POD, SOD and CAT activities were all highly significantly positively correlated with total phenolic content (*P* < 0.01), and CAT was highly significantly positively correlated with tannin and total flavonoid contents (*P* < 0.01), indicating that elevated defense enzyme activities are closely coupled with secondary metabolite synthesis and accumulation, possibly jointly enhancing resistance via activation of the phenylpropanoid pathway. Total phenolics, tannins and total flavonoids were all highly significantly positively correlated with one another (*P* < 0.01), indicating synchronous and synergistic accumulation of these resistance-related compounds, jointly constituting an important component of chemical defense. POD, SOD and CAT activities were all highly significantly negatively correlated with amino acid content (*P* < 0.01), and tannin content was highly significantly negatively correlated with both amino acid and soluble sugar contents (*P* < 0.01), suggesting that under feeding stress part of the nitrogen and carbohydrate resources is reallocated toward defensive secondary metabolite or stress protein synthesis, relatively reducing nutrient contents. In summary, after feeding–infestation the defense enzyme activities and secondary metabolites increased significantly and synergistically, while defensive substance accumulation was significantly negatively correlated with certain basic nutrients, reflecting pronounced resource reallocation and metabolic trade-off.

**Figure 5.**
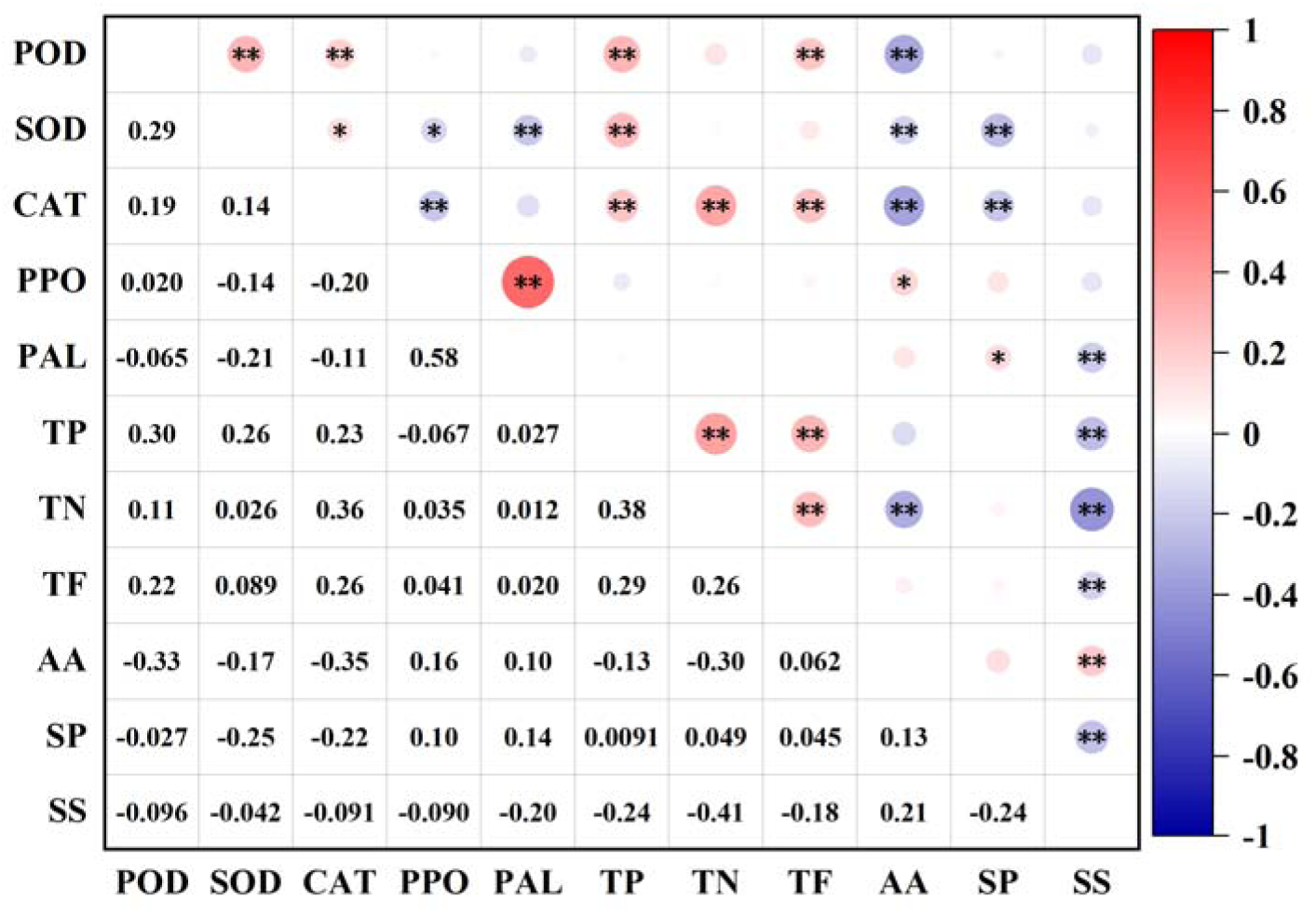
Correlation Analysis of Resistance Physiological Indicators of Clones after Infestation by *D. superans* Note: * indicates significant difference (P < 0.05), ** indicates highly significant difference (P < 0.01). TP, total phenols; TN, tannins; TF, total flavonoids; AA, amino acids; SP, soluble protein; SS, soluble sugar.

Cluster analysis after feeding (Fig. 6) separated the 46 clones at a genetic distance of 450 into three categories: Category I (41 clones) with relatively higher PPO and PAL activities and tannin, total flavonoid, amino acid and soluble protein contents; Category II (4 clones) with relatively higher POD, SOD, CAT and PPO activities and total phenolic, tannin, total flavonoid and soluble sugar contents; and Category III (one clone) with relatively higher CAT activity and total phenolic, amino acid and soluble sugar contents. The clones thus exhibited a pronounced induced resistance response through synergistic enhancement of defense enzymes and secondary metabolites accompanied by nutrient reallocation, providing a physiological basis for elucidating resistance mechanisms at the metabolic level.

**Figure 6.**
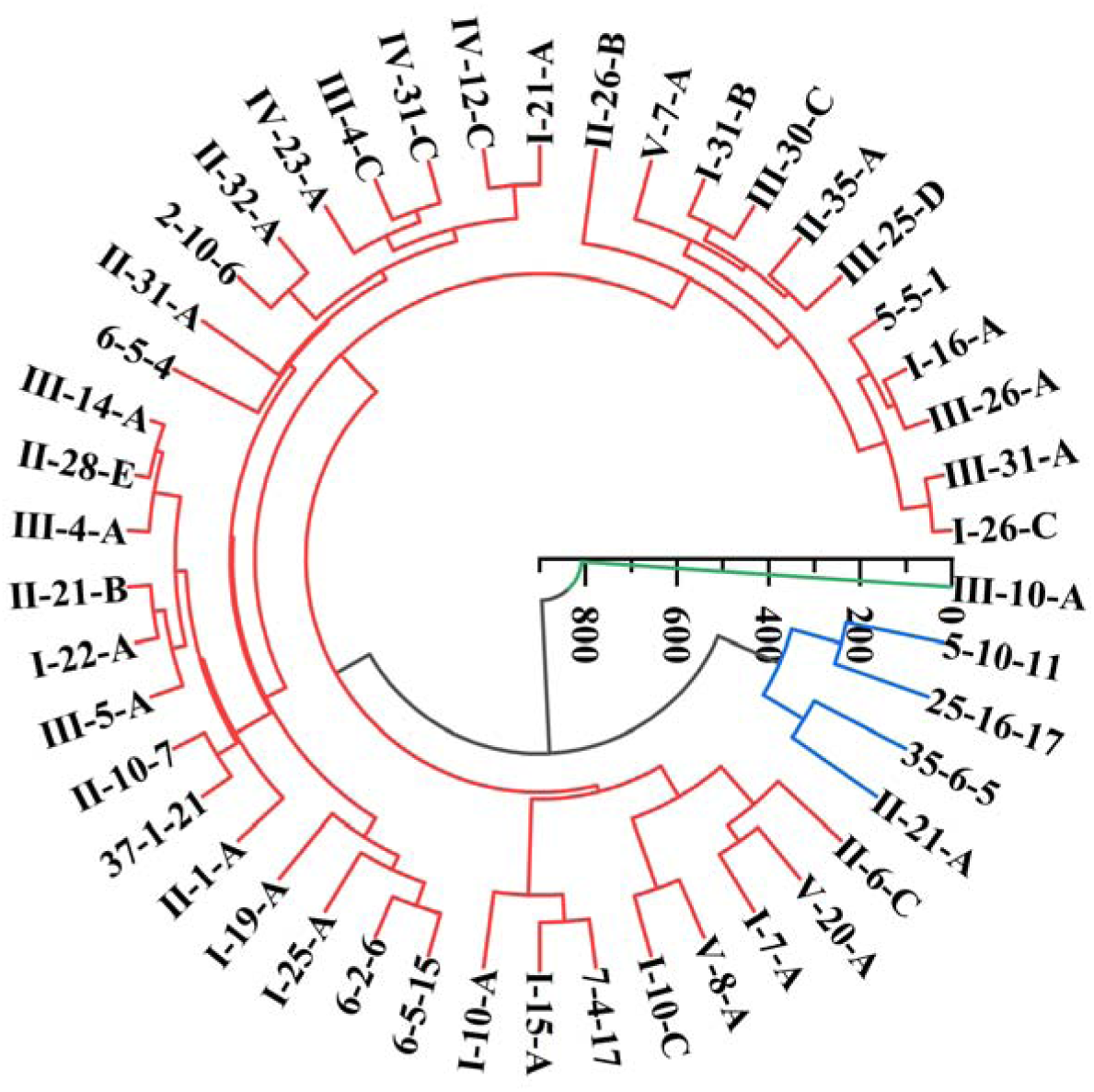
Cluster Analysis of Clones after Infestation by *D. superans*

### 3.4 Comprehensive Evaluation by the Membership Function Method

#### 3.4.1 Comprehensive Evaluation of the Clones Based on Insect Weight, Frass Weight and Defoliation Rate

Using the membership function method, the insect resistance of the 46 hybrid larch clones was comprehensively evaluated with insect weight, frass weight and clone defoliation rate as evaluation indices. The comprehensive evaluation scores and rankings of all clones are shown in Table S4. The D values of -30-C, -35-A, -21-A, -7-A and -26-A were relatively high compared with those of the other clones, being 0.960, 0.960, 0.952, 0.951 and 0.939, respectively, indicating their superior insect resistance and providing a theoretical basis for the subsequent screening of insect-resistant materials.

#### 3.4.2 Comprehensive Evaluation of the Clones Based on Indices before Feeding–infestation

Based on the resistance-related physiological indices of the clones before feeding–infestation by the *D. superans*, a comprehensive evaluation of the 46 hybrid larch clones was performed using the membership function method. The comprehensive evaluation scores and rankings of all clones are shown in Table S5. The D values of -28-E, 6-5-15, -20-A, -1-A and -5-A were relatively high, being 0.761, 0.740, 0.709, 0.707 and 0.635, respectively, indicating that these five clones possessed relatively high constitutive resistance.

#### 3.4.3 Comprehensive Evaluation of the Clones Based on Indices after Feeding–infestation

Based on the resistance-related physiological indices of the clones after feeding–infestation by the *D. superans*, a comprehensive evaluation of the 46 hybrid larch clones was performed using the membership function method. The comprehensive evaluation scores and rankings of all clones are shown in Table S6. The comprehensive evaluation scores of -28-E, -20-A, -1-A, 6-5-15 and -5-A were the highest, being 0.688, 0.678, 0.657, 0.648 and 0.582, respectively, indicating that these five clones possessed relatively high induced resistance.

Overlap analysis across the three perspectives (Table 7) showed no overlap between clones screened as superior in insect resistance and those superior in constitutive or induced resistance, whereas clones superior in constitutive and induced resistance overlapped to a high degree. Insect resistance assessed by feeding consequences and intrinsic resistance assessed by physiological indices thus reflect different facets of resistance, so evaluation from any single perspective is insufficient to fully characterize the true resistance level, and multi-perspective evaluation is more reliable.

**Table 7.**
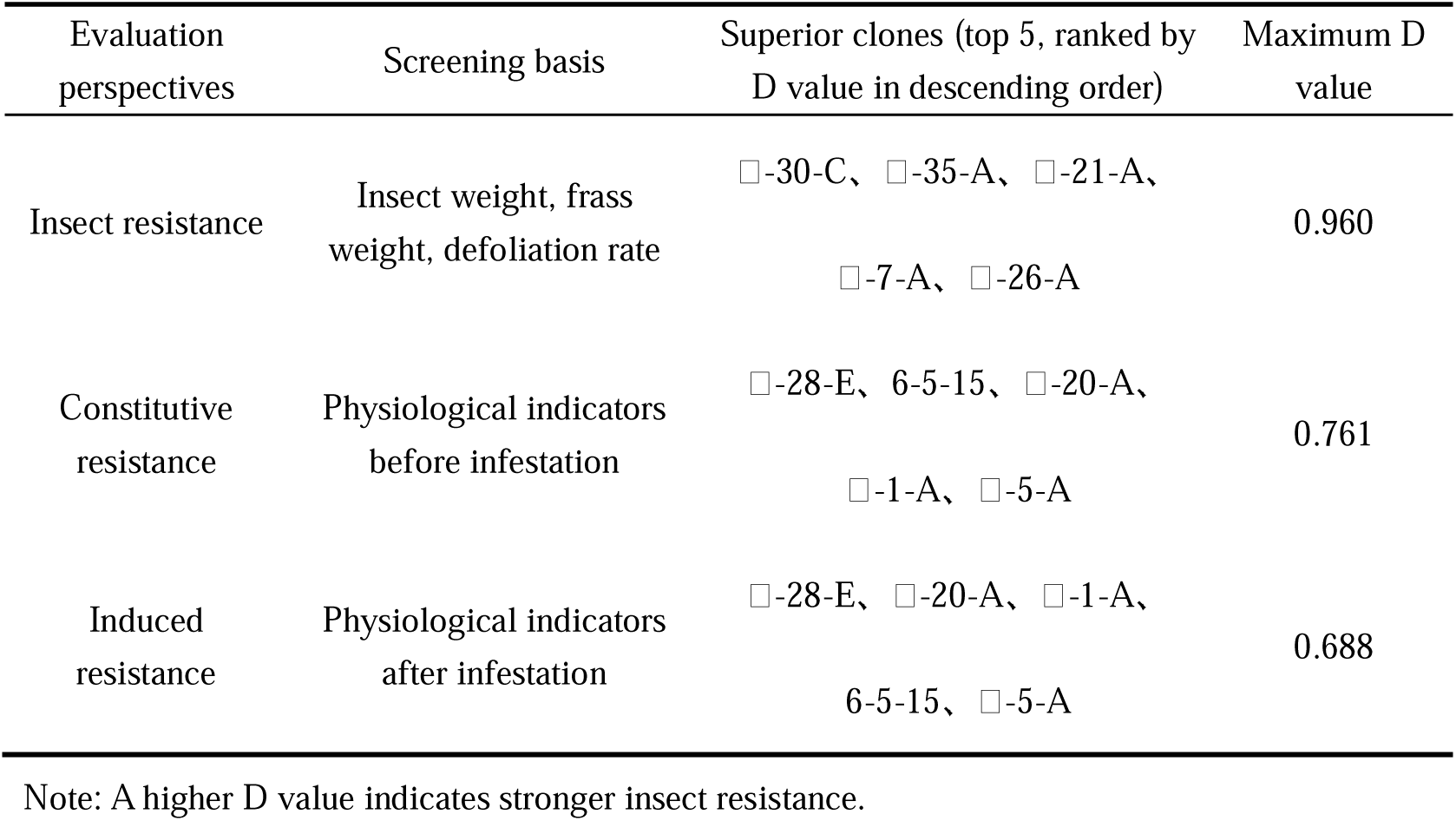
Comparison of Optimal Screening Results by Subordinate Function Comprehensive Evaluation from Three Evaluation Perspectives.

### 3.5 Metabolomic Analysis

Because the superior materials screened from the three perspectives were not fully consistent, high- and low-resistance clones from each perspective were subjected to widely targeted LC-MS metabolomics (6 groups, 18 samples, 3 biological replicates each) to elucidate the principal defensive substances and mechanisms by which hybrid larch resists *D. superans* feeding. The comparisons were: insect resistance (HIR, -30-C vs LIR, -14-A); constitutive resistance (HPR, -28-E vs LPR, -10-A); and induced resistance (HCR, -28-E vs LCR, -31-A).

#### 3.5.1 Differential Metabolite Analysis

Widely targeted LC-MS metabolomic analysis was performed on the above six groups comprising 18 plant samples in total, and a total of 3,577 metabolites were identified. In terms of chemical classification (Fig. 7), the dominant metabolites were terpenoids: diterpenoids accounted for 26.5%, sesquiterpenoids 22.6%, triterpenoids 15.1%, monoterpenoids 10.5%, steroids 9.9%, meroterpenoids 4.8%, carotenoids (C) 3.7%, apocarotenoids 3.0%, and sesterterpenoids 0.4%, while pseudoalkaloids (transamidation), polyprenols, polyethers, pseudoalkaloids and tryptophan alkaloids each accounted for 0.1%. In total, terpenoids and their derivatives accounted for more than 90% of the identified metabolites, indicating that terpenoid metabolites constitute the principal component of the needle metabolite repertoire of hybrid larch and represent its potential chemical defense basis.

**Figure 7.**
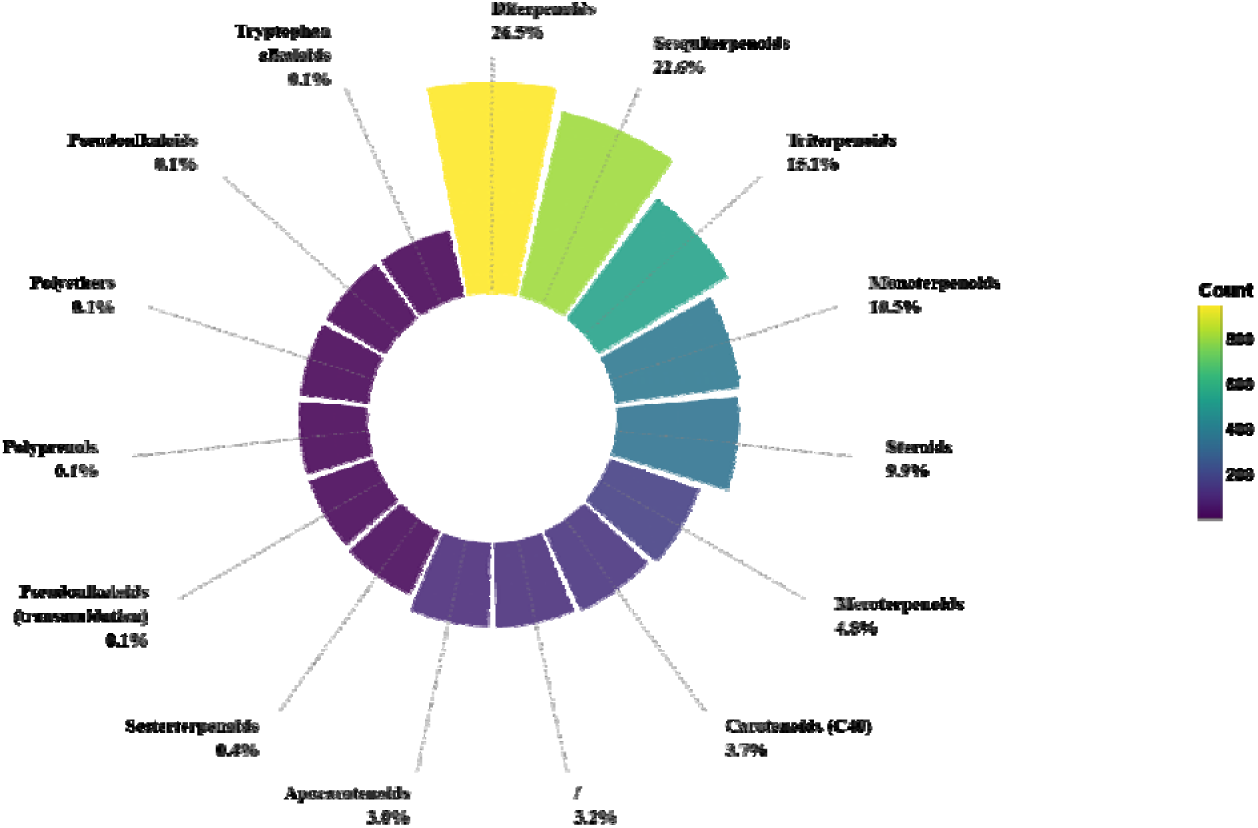
Classification and Proportion of Differentially Expressed Metabolites in Hybrid Larch Clones

Differential metabolites were screened using the criteria of |log Fold Change| > 1, PADJ < 0.05 and VIP > 1 in the OPLS-DA model, and the numbers of differential metabolites in each comparison group were displayed by Venn diagrams and volcano plots (Fig. 8). As shown in Fig. 8A, the three comparison groups (HIR vs LIR, HPR vs LPR and HCR vs LCR) shared 83 differential metabolites, whereas the numbers of group-specific differential metabolites were 503, 547 and 543, respectively, indicating that the metabolic responses of the resistance materials screened from different evaluation perspectives to *D. superans* feeding share both commonalities and differences. As shown in Figs. 8B–8D, 1,010 differential metabolites were identified in HIR vs LIR, of which 541 were up-regulated and 469 down-regulated; 1,086 differential metabolites were identified in HPR vs LPR, of which 513 were up-regulated and 573 down-regulated; and 922 differential metabolites were identified in HCR vs LCR, of which 616 were up-regulated and 306 down-regulated. Notably, in HCR vs LCR the number of up-regulated metabolites was markedly greater than that of down-regulated metabolites, suggesting that, after feeding stress, the induced-resistance materials predominantly exhibited positive accumulation of metabolite synthesis, which is consistent with the enhanced resistance physiology observed after feeding–infestation.

**Figure 8.**
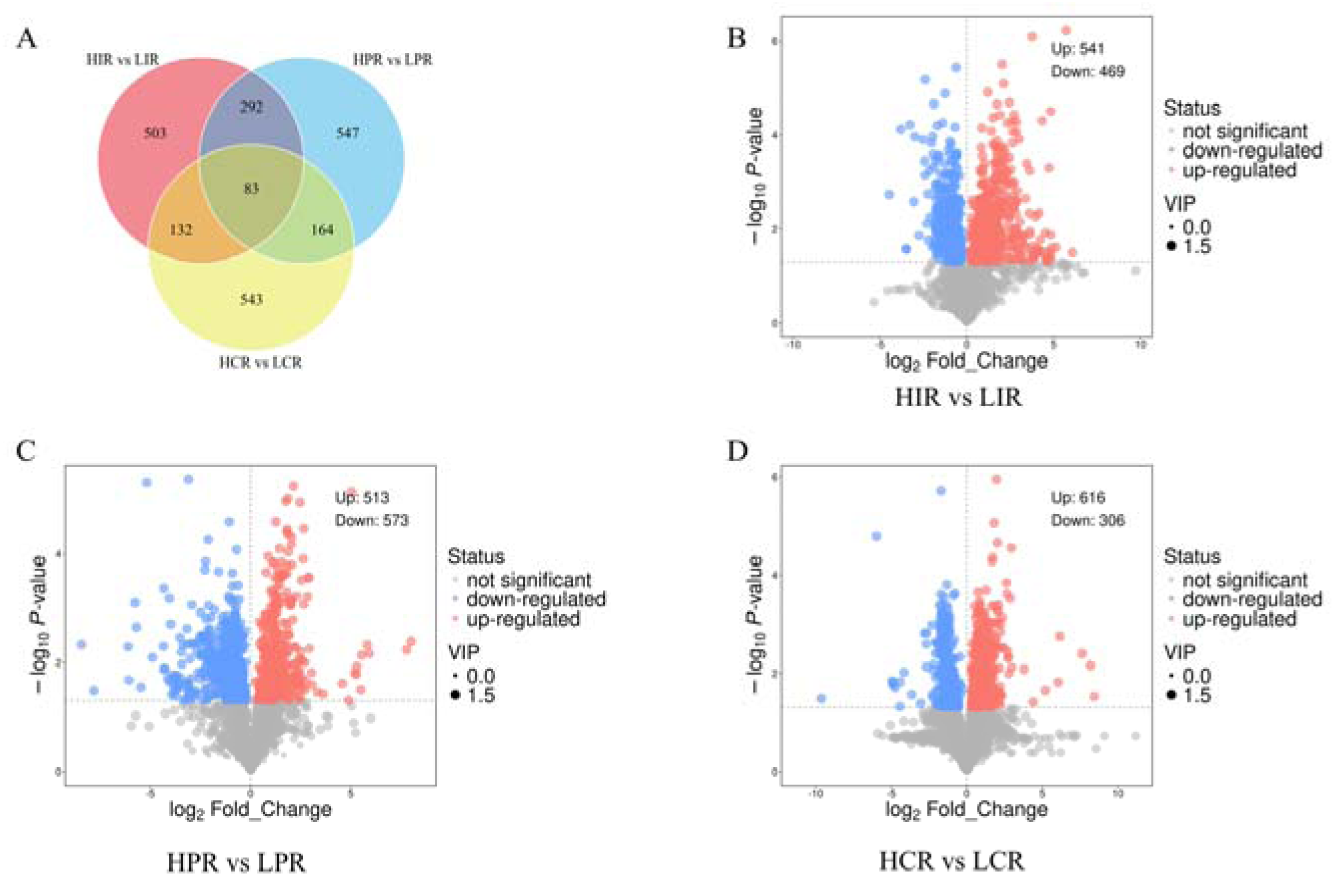
Venn Diagram (A) and Volcano Plots (B: HIR vs LIR; C: HPR vs LPR; D: HCR vs LCR) of Differentially Expressed Metabolites among Clones with Different Resistance Levels

#### 3.5.2 Principal Component Analysis

To further compare the overall metabolic differences among hybrid larch clones of different resistance categories under *D. superans* feeding stress, principal component analysis (PCA) was performed separately on the metabolite data of the three comparison groups (HPR vs LPR, HCR vs LCR and HIR vs LIR), and the results are shown in Fig. 9. As shown in Fig. 9A, the HPR and LPR samples exhibited a clear separation trend along the PC1 axis (explained variance 64.1%), with a PC2 explained variance of 13.2%; the two groups clustered on the positive and negative halves of PC1, respectively, indicating that the metabolite composition of the high- and low-resistance clones already differed significantly before feeding–infestation. In Fig. 9B, the HCR and LCR samples were separated along the PC1 axis (explained variance 48.2%), with a PC2 explained variance of 23.6%, and the two categories of clones formed distinct clusters. In Fig. 9C, the HIR and LIR samples also showed clear discrimination along the PC1 axis (explained variance 63.0%). The cumulative variance contribution rates of PC1 and PC2 in the three comparison groups were 77.3%, 71.8% and 76.2%, respectively, all of which adequately reflected the characteristics of the original data. Overall, the samples of clones of different resistance categories were effectively separated in the PCA score plots, indicating good inter-group discrimination and reproducibility of their metabolite composition, thereby laying a foundation for the subsequent analysis of differential metabolites and metabolic pathways.

**Figure 9.**
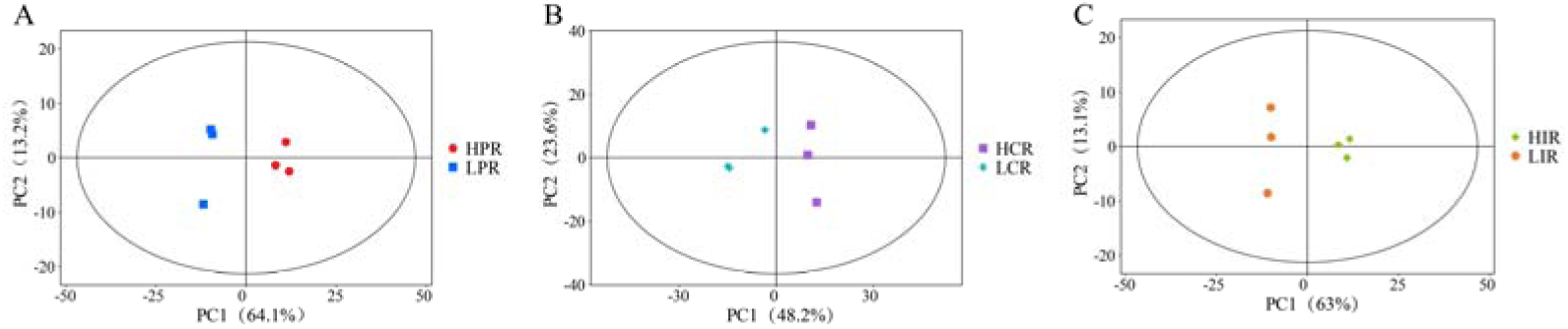
Principal Component Analysis of Metabolites in Clones with Different Resistance Levels before and after Infestation by *D. superans*

#### 3.5.3 KEGG Enrichment Pathway Analysis

To interpret the biological processes in which the differential metabolites participate, KEGG pathway enrichment analysis was further performed on the differential metabolites of each comparison group, and the results are shown in Fig. 10. As shown in Fig. 10A (HPR vs LPR), the differential metabolites were mainly enriched in pathways such as metabolic pathways, biosynthesis of secondary metabolites, monoterpenoid biosynthesis, biosynthesis of various plant secondary metabolites, diterpenoid biosynthesis, and phenylpropanoid biosynthesis. As shown in Fig. 10B (HIR vs LIR), in addition to the aforementioned shared pathways, the differential metabolites were also significantly enriched in pathways such as steroid biosynthesis, glycerolipid metabolism and tyrosine metabolism. As shown in Fig. 10C (HCR vs LCR), the carotenoid biosynthesis pathway was markedly enriched. Integrating the results of the three comparison groups, “metabolic pathways” and “biosynthesis of secondary metabolites” constitute the basal pathways commonly enriched by the differential metabolites in all comparison groups, whereas the biosynthetic pathways of terpenoid and phenolic secondary metabolites—such as monoterpenoids, diterpenoids, sesquiterpenoids and phenylpropanoids—were consistently enriched across multiple comparison groups. This indicates that, in response to feeding stress by the *D. superans*, the hybrid larch clones mainly synthesize defensive metabolites by regulating the terpenoid and phenylpropanoid secondary metabolic pathways.

**Figure 10.**
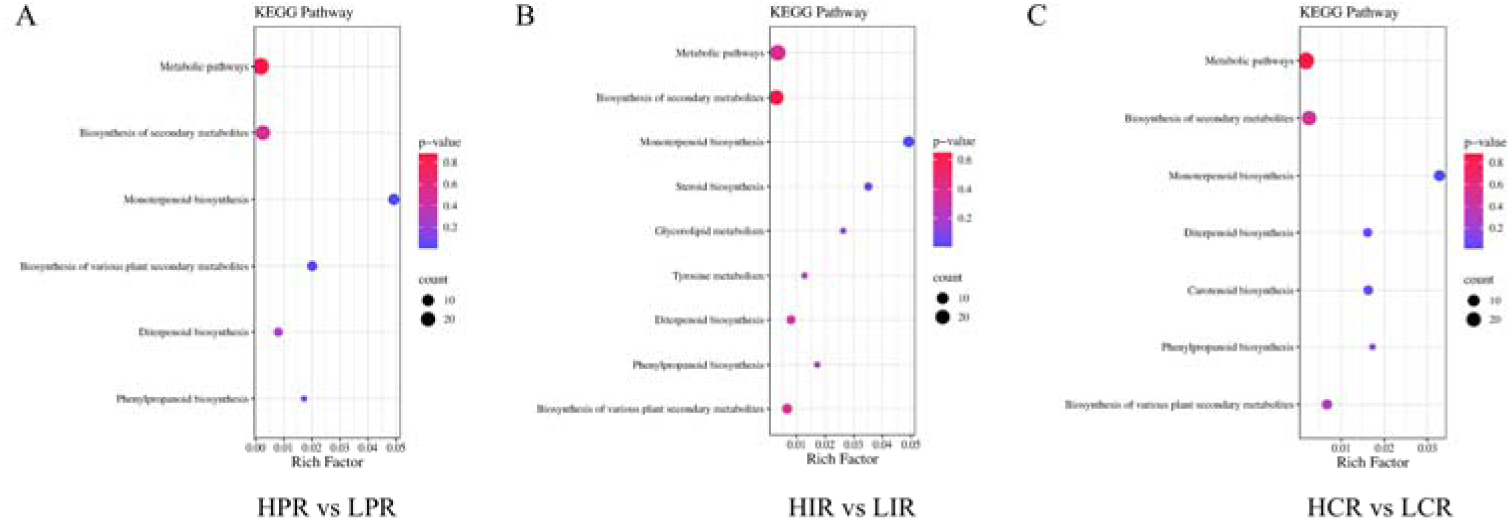
KEGG Enrichment Pathway Analysis of Differentially Expressed Metabolites in Clones with Different Resistance Levels before and after Infestation by *D. superans*

**Figure 11.**
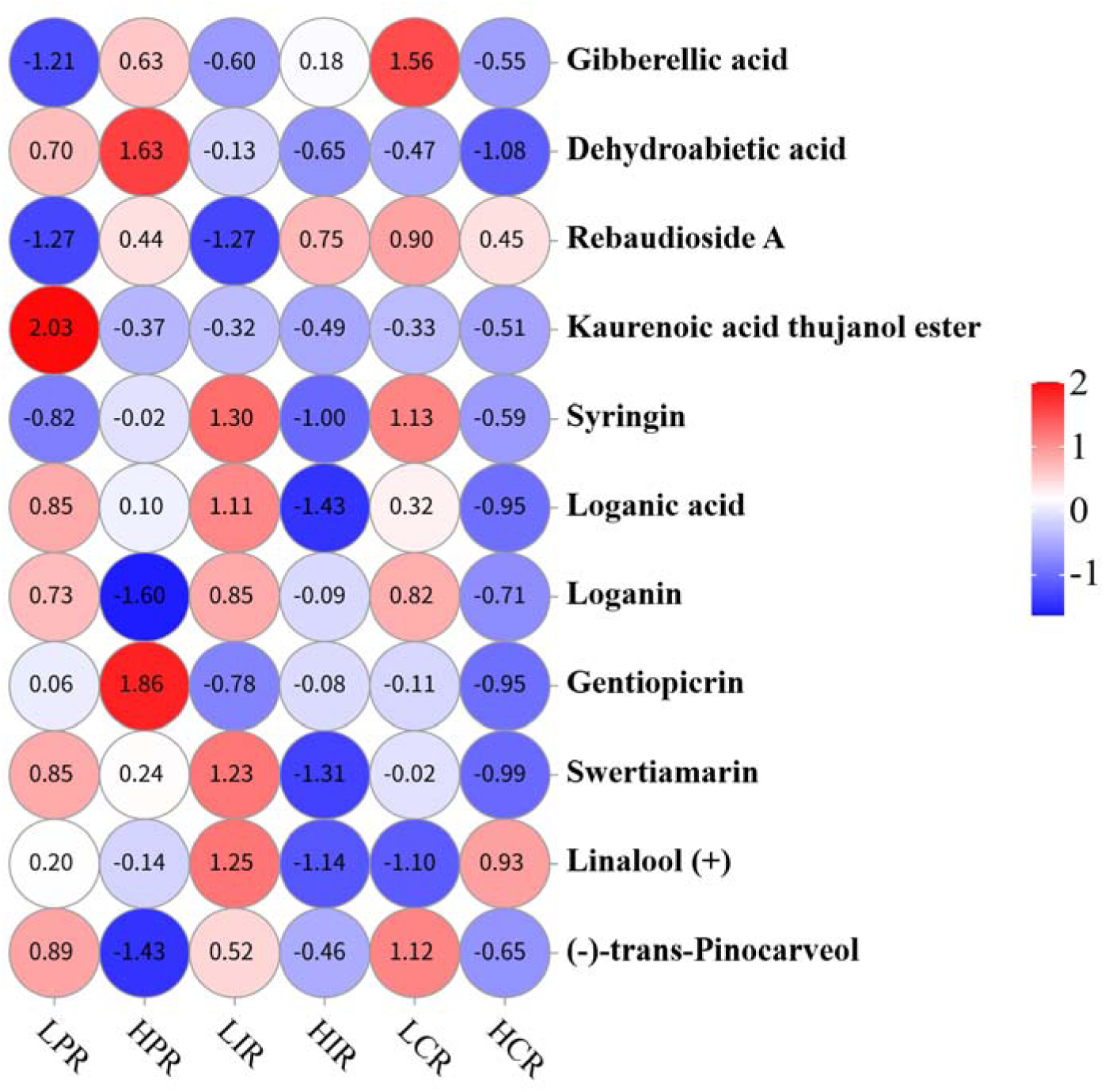
Heatmap Analysis of Key Differentially Expressed Metabolites

Further analysis of specific differential metabolites revealed that, in HPR vs LPR, terpenoid and phenolic compounds such as gibberellic acid, syringin, gentiopicrin and dehydroabietic acid were significantly up-regulated in superior clones; in HIR vs LIR, diterpenoids such as gibberellic acid and rebaudioside A were significantly accumulated; and in HCR vs LCR, many monoterpene glycosides, diterpenoids and sesquiterpenoids, such as the iridoid glycosides loganin, loganic acid and swertiamarin, were significantly up-regulated in high-resistance materials after feeding. Among these, gibberellic acid, dehydroabietic acid and kaurenoic acid are typical diterpenoid and resin acid compounds constituting important chemical defense substances of conifers against insect feeding, whereas syringin is a key phenolic product of the phenylpropanoid pathway. Consistent with the physiological results, after feeding–infestation the defense enzyme activities and secondary metabolite contents increased significantly and synergistically and were significantly negatively correlated with nutrients such as amino acids and soluble sugars. The metabolomic results further showed that, under feeding stress, resistant materials were significantly enriched in terpenoid and phenylpropanoid pathways with terpenoids and phenolics as predominant metabolites, highly consistent with the physiological synergy. It can thus be concluded that, in resisting *D. superans* feeding, hybrid larch clones may primarily rely on terpenoid and phenolic compounds, jointly constituting a chemical defense network through synergistic regulation of the terpenoid and phenylpropanoid pathways, and enhance needle resistance through resource reallocation between primary and secondary metabolism, also explaining at the metabolic level the differences among materials screened from different perspectives.

## Discussion

*D. superans* is a major defoliating pest of larch plantations, and its feeding directly affects tree growth and wood quality^26^. Breeding and screening insect-resistant superior larch materials is the most economical and fundamental control approach. In this study, insect weight, frass weight and defoliation rate all exhibited highly significant differences after field feeding (P < 0.01) and were significantly or highly significantly positively correlated, indicating abundant genetic variation and that these indices jointly characterize resistance from feeding amount, digestive metabolism and damage consequences. Notably, superior materials screened from a single perspective were not fully consistent: insect-resistant superior materials based on the three damage indices (e.g., -30-C, -35-A), constitutive-resistance materials based on pre-feeding indices (e.g., -28-E, 6-5-15) and induced-resistance materials based on post-feeding indices (e.g., -28-E, -20-A) showed partial but not identical overlap. This suggests that insect resistance is a complex trait regulated by multiple genes and pathways, and that evaluation based on a single hierarchical level or time point cannot fully reflect the true resistance level of a material^27^.

The resistance of plants to defoliating pests is the result of the synergistic action of constitutive resistance and induced resistance, which are driven by distinct physiological and metabolic mechanisms^28,29^. In this study, before feeding–infestation, POD and SOD activities in the clones were highly significantly positively correlated, indicating their synergistic action in scavenging ROS, whereas SOD activity was highly significantly negatively correlated with PPO and PAL activities^30,31^, possibly reflecting the functional differentiation and substrate competition among defense enzyme systems prior to feeding stress^32,33^. After feeding–infestation, POD, SOD and CAT activities, as well as PPO and PAL activities, were all highly significantly positively correlated, and the defense enzyme activities increased in synchrony with the contents of secondary metabolites such as total phenolics, tannins and total flavonoids, indicating that, after damage, the enzymatic defense and secondary metabolite defense of the clones were synergistically activated and the induced resistance response was rapidly amplified^34^. Previous studies have shown that defense enzymes such as POD, PPO and PAL play key roles in cell wall reinforcement and barrier formation by participating in the oxidative cross-linking of phenolics and lignin deposition^35–37^, whereas CAT and SOD directly scavenge ROS to alleviate oxidative damage^38–40^. Therefore, before feeding, the clones rely more on pre-existing constitutive resistance, whereas after feeding they rapidly activate induced resistance centered on enzymatic defense and secondary metabolite synthesis. This dynamic transition determines the ultimate capacity of the clones to resist the *D. superans*, and also corroborates, at the physiological level, the necessity of multi-perspective evaluation.

Secondary metabolites constitute an important chemical basis for plant resistance to pest feeding, among which terpenoids and phenolic compounds are the core components of the chemical defense of conifers^41–43^. In this study, a total of 3,577 metabolites were identified by metabolomic analysis, with terpenoids and their derivatives accounting for more than 90%, which is consistent with the characteristic richness of terpenoid secondary metabolites in conifers and further indicates that terpenoid metabolism constitutes the metabolic basis of the chemical defense of hybrid larch^43^. Regarding the differential metabolites, diterpenoid and resin acid compounds such as gibberellic acid, dehydroabietic acid and rebaudioside A were significantly enriched in the high-resistance materials, whereas phenylpropanoids and iridoid glycosides, such as syringin, loganic acid, loganin and gentiopicrin, were present at higher levels in the low-resistance materials. Previous studies have confirmed that diterpene resin acids, represented by abietic acid, are important defensive substances of conifers against both bark-boring and defoliating pests, exerting direct antifeedant and toxic effects^44–46^; they can inhibit insect feeding, growth and development by interfering with the insect nervous or digestive system. Iridoid glycosides (such as loganin, loganic acid and their derivatives) likewise exhibit pronounced insecticidal activity and can inhibit the feeding and growth of various pests^47,48^. In addition, the monoterpene volatiles released by larch, such as α-pinene and myrcene, exert an attractive effect on *D. superans*^49^, whereas camphene exhibits a repellent effect.^50^ This indicates that monoterpenes not only participate directly in defense but may also influence pest host-selection behavior through volatile signals^51^.

A point worthy of in-depth discussion is that the significant enrichment of gibberellic acid in the high-resistance materials appears to be in tension with the classical view that growth-promoting hormones usually weaken defense. On the one hand, gibberellic acid, as a diterpenoid phytohormone, is itself a product of the terpenoid metabolic pathway^52^; the enrichment of gibberellic acid and its intermediates in the high-resistance materials may reflect the overall up-regulation of the terpenoid metabolic pathway in resistant materials, serving as a metabolic signature of the activation of the terpenoid defense network^53^. On the other hand, previous studies have shown that the gibberellic acid signaling pathway cross-regulates and antagonizes the jasmonic acid signaling pathway^54,55^, and that exogenous gibberellic acid inhibits JA-mediated defense responses, thereby reducing plant resistance to herbivorous insects^56,57^. It can thus be inferred that the gibberellic acid enriched in the high-resistance materials in this study may not exert a direct insecticidal role, but rather serve as a marker of active terpenoid metabolism, or participate in regulating the resource allocation that balances growth and defense. This phenomenon also suggests that insect resistance is not determined by a single compound, but should be understood within the framework of a multi-metabolite interaction network.

Integrating the differential metabolite and KEGG results, in response to feeding the clones mainly synthesize defensive metabolites through synergistic regulation of the terpenoid and phenylpropanoid pathways, together constituting the “terpenoid–phenolic” chemical defense network (Fig. 12). Terpenoids mainly exert direct antifeedant and toxic effects, interfering with pest feeding, digestion and growth and development^19^. Phenolics, in turn, not only participate in cell wall reinforcement as lignin synthesis precursors to form a physical barrier, but also exhibit antioxidant, insecticidal and antimicrobial activities^58^, and can further strengthen defense as precursors of phytoalexins^59,60^. This dual synergistic mechanism was reflected in the resistant materials screened from different evaluation perspectives: KEGG enrichment analysis showed that all comparison groups were significantly enriched in pathways such as monoterpenoid biosynthesis, diterpenoid biosynthesis and phenylpropanoid biosynthesis, and the direction of the differences in the metabolomic results corroborated the physiological findings of elevated defense enzyme activities and increased contents of total phenolics, tannins and total flavonoids after feeding–infestation, indicating that the metabolic defense and physiological defense of the clones are highly coupled in response to feeding stress.

**Figure 12.**
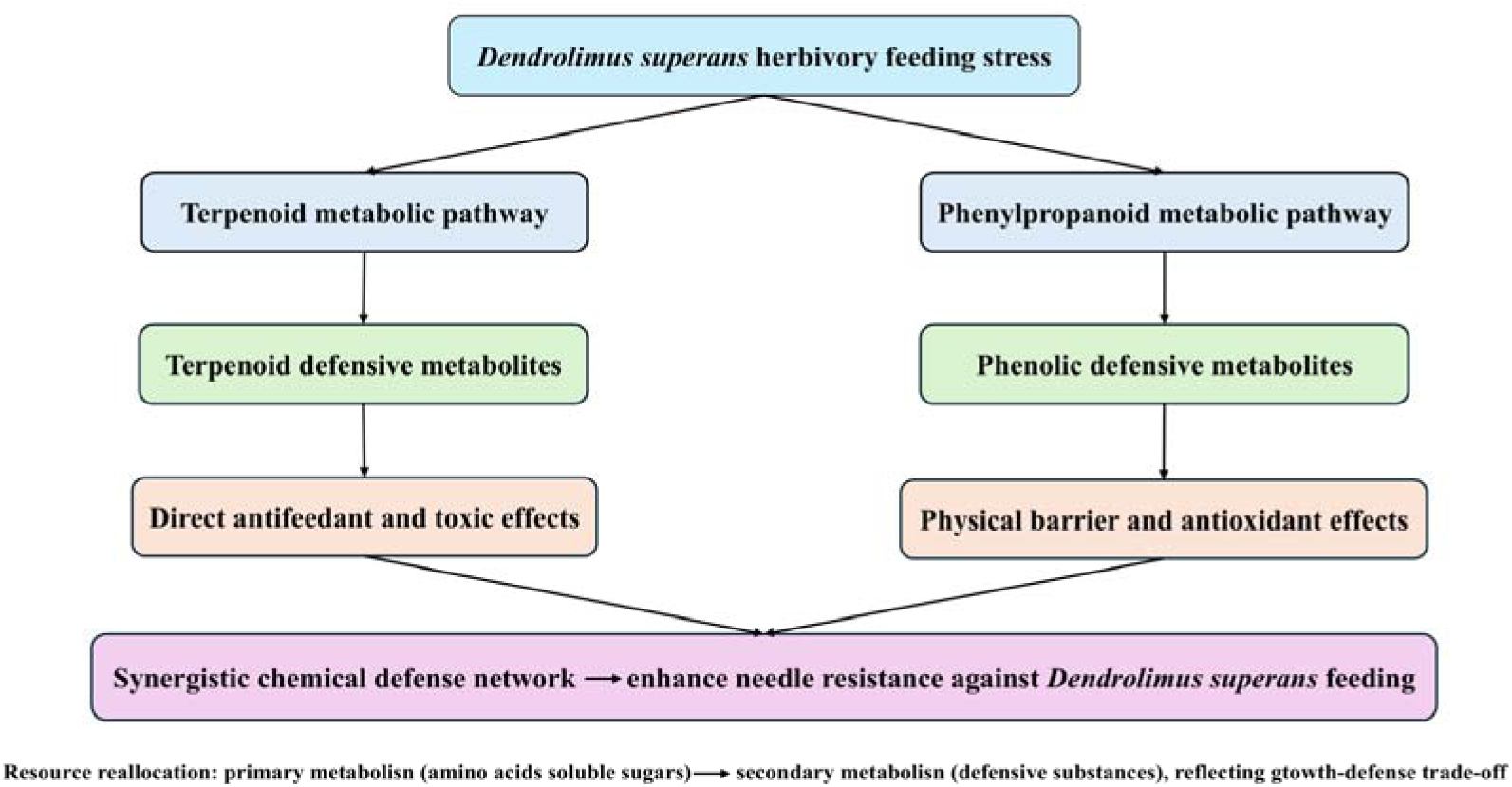
Schematic Diagram of Defense Network

When subjected to feeding stress by pests, plants often face a resource trade-off between growth and defense^61,62^. The correlation analysis in this study showed that secondary metabolites such as total phenolics, tannins and total flavonoids were significantly or highly significantly positively correlated with one another, but highly significantly negatively correlated with the contents of nutrients such as amino acids and soluble sugars. This suggests that, after damage, the clones reallocate part of their carbon and nitrogen resources from primary metabolism toward the synthesis of defensive secondary metabolites. This resource reallocation is consistent with the growth–defense trade-off theory^63–65^. According to the carbon–nutrient balance hypothesis and the optimal defense hypothesis, investment in defense consumes resources that could otherwise be used for growth^66–68^; therefore, insect resistance breeding needs to strike a balance between resistance and growth. It should be noted that this study did not directly measure the growth indices of the clones; the analysis of the growth–defense trade-off herein is an indirect inference based on the correlations among resistance, defensive substances and nutrients, and the quantitative relationship between growth and resistance still requires verification through the subsequent measurement of growth traits. For hybrid larch, the high-resistance materials in this study were able to maintain a relatively high level of defensive metabolism after feeding damage, and some materials (e.g., -28-E) performed well across all three perspectives of insect resistance, constitutive resistance and induced resistance, implying that genotypes balancing both resistance and growth may exist and could serve as priority materials for insect resistance breeding. Meanwhile, integrating the results from the phenotypic, physiological and metabolic levels, it can be seen that the resistance of hybrid larch clones to the *D. superans* is a complex system jointly constituted by larval developmental phenotype, the defense enzyme system, the accumulation of secondary metabolites, and the terpenoid–phenolic metabolic network. Nevertheless, this study revealed the resistance differences only at the metabolomic and physiological-biochemical levels; the molecular regulatory mechanisms of the key defensive metabolites, the verification of their biological activity through artificial feeding assays, and their stability in long-term field pest management still warrant further investigation via transcriptomics, metabolite feeding experiments, and multi-site, multi-year validation.

## Conclusion

Using 46 hybrid larch clones, this study comprehensively evaluated insect resistance with the membership function method through field feeding–infestation experiments from three perspectives—insect weight, frass weight and defoliation rate; pre-infestation resistance-related physiological indices; and post-infestation resistance-related physiological indices—and conducted widely targeted LC-MS metabolomics on representative materials. The main conclusions are as follows:

1. Insect weight change, frass weight and defoliation rate all exhibited highly significant differences among clones (*P* < 0.01) and were significantly or highly significantly positively correlated, indicating abundant genetic variation in resistance to *D. superans*. Based on the comprehensive evaluation with the membership function method, clones such as -30-C, -35-A and -21-A showed strong insect resistance; clones such as -28-E, 6-5-15 and -20-A showed high constitutive resistance; and clones such as -28-E, -20-A and -1-A showed high induced resistance.
2. Before feeding, enzymatic defense, secondary metabolites and nutrients were highly interrelated: POD and SOD changed synergistically, and total phenolics, tannins and total flavonoids accumulated synergistically while showing significant negative correlations with nutrients such as amino acids. After feeding, defense enzyme activities and secondary metabolite contents increased significantly and synergistically, indicating that the clones enhance resistance through the synergistic action of enzymatic defense and secondary metabolite defense.
3. A total of 3,577 metabolites were identified, predominantly terpenoids (>90%). The differential metabolites of the three comparison groups were dominated overwhelmingly by terpenoids and commonly enriched in metabolic pathways, biosynthesis of secondary metabolites, monoterpenoid and diterpenoid biosynthesis, and phenylpropanoid biosynthesis. In resistant materials, diterpenoid and resin acid compounds such as gibberellic acid, dehydroabietic acid and kaurenoic acid, together with phenolic compounds such as syringin, were significantly up-regulated (gibberellic acid, as a diterpenoid phytohormone, may be associated with overall terpenoid pathway activation rather than acting as a direct insecticidal factor, warranting further verification). The clones thus constitute a chemical defense network dominated by terpenoid and phenolic compounds through synergistic regulation of the terpenoid and phenylpropanoid pathways, enhancing needle resistance to *D. superans* feeding.

In summary, this study revealed the resistance mechanism of hybrid larch clones to *D. superans* at the phenotypic, physiological and metabolic levels. The superior materials screened for insect resistance, constitutive resistance and induced resistance provide a material basis for the selection and breeding of insect-resistant superior hybrid larch clones, and the terpenoid–phenolic synergistic defense network revealed herein serves as a theoretical reference for insect resistance breeding in conifers.

## Author contributions

This project was conceived by LCM and BH, with YPK as the main writer. LML, X LH, and LJ provided substantial assistance during field experiments. All authors edited and approved the manuscript.

## Acknowledgments

This study was supported by the National Science and Technology Major Project of the Ministry of Science and Technology (grant no. ZKJB230100017).

